# TGAC Browser: An open-source genome browser for non-model organisms

**DOI:** 10.1101/677658

**Authors:** Anil S. Thanki, Xingdong Bian, Robert P. Davey

## Abstract

Genome browsers play a vital role to provide visualisation for genomic data. It is often the case that bespoke genome browser customisations are required between different research groups, with an obvious necessity to update, upgrade and tailor tracks and features on a potentially frequent basis. However, most of the current genome browsers require highly curated data held in public repositories. Besides, these genome browsers often rely on particular dependencies, where writing plug-in or modifying existing code can be troublesome and resource expensive.

We present TGAC Browser, a new open-source web-based genome browser designed to overcome shortcomings in available approaches. It uses a locally installed Ensembl Core Database schema and is also able to visualise data from well-known NGS data formats. We also added simple analysis functionality to perform BLAST searches within TGAC Browser. TGAC Browser also allows uploading your genomic data. TGAC Browser is an open-source, easy to set up, and user-friendly genome browser with minimal, lightweight configuration details.

## Introduction

Genome browsers (1–3) typically present spatial relationships between different pieces of biological information by providing graphical visualisations of the genomic data. Despite advances in data production and analysis methods, genome browsers play an important role in examining data to explore the results of new analysis and generating hypotheses (4). The principal function of the genome browser is to aggregate different types of genomic annotation data together and integrate them into an abstract graphical view (5). It allows researchers to visualise and explore predicted genes, transcripts, gene expression, variation, comparative analysis, and alignments. Because of so many of these reasons to use genome browser, many software has been developed which are widely used and essential, for example, Ensembl genome browser (3), GBrowse (1), JBrowse (6), and IGV (7).

In general, genome browsers can be divided into two categories: standalone browsers and web-based browsers. Standalone browsers are used on a local computer, which tends to focus on heavyweight applications to run with a large dataset. While web-based browsers are generally installed on host institutional server and can be used over the internet. Here we are focusing on a web-based genome browser, which is more popular due to its flexible accessibility and performance.

Many of the available web-based genome browsers require heavily curated data in public repositories as well as in a specific format supported by the particular browser. With the democratisation of sequencing technologies, smaller research labs are generating an increasing amount of sequencing data and performing analyses, especially for non-model organisms. Biological analyses can be performed in many alternative ways providing results in various formats; thus it can be a tedious process to convert data in order for it to be supported by a particular browser and data needs to be curated before making them available from a public repository. Among various available genomic formats, the Ensembl database system (8) is standard format containing various genomic annotation, and it is widely accepted in both companies as well as academic sites and also provides a framework to load any standard NGS formatted data into Ensembl databases.

To better simplify genomic data visualisation, we present the TGAC Browser, an open-source genome browser. It retrieves and visualises data directly from a local instance of the Ensembl core database (8) as well as well-known NGS data formats. TGAC Browser can also perform BLAST (Basic Local Alignment Search Tool) (9) analysis within.

## Materials and Methods

TGAC Browser is designed with a typical server-client architecture (see Figure 1) to utilise the server for data retrieval and use clients’ computational resources to generate the visualisation. This approach provides a consistent experience to users when the TGAC Browser is being used by multiple users simultaneously. Client and server transfer data asynchronously using Ajax (Asynchronous JavaScript And XML).

**Fig. 1.**
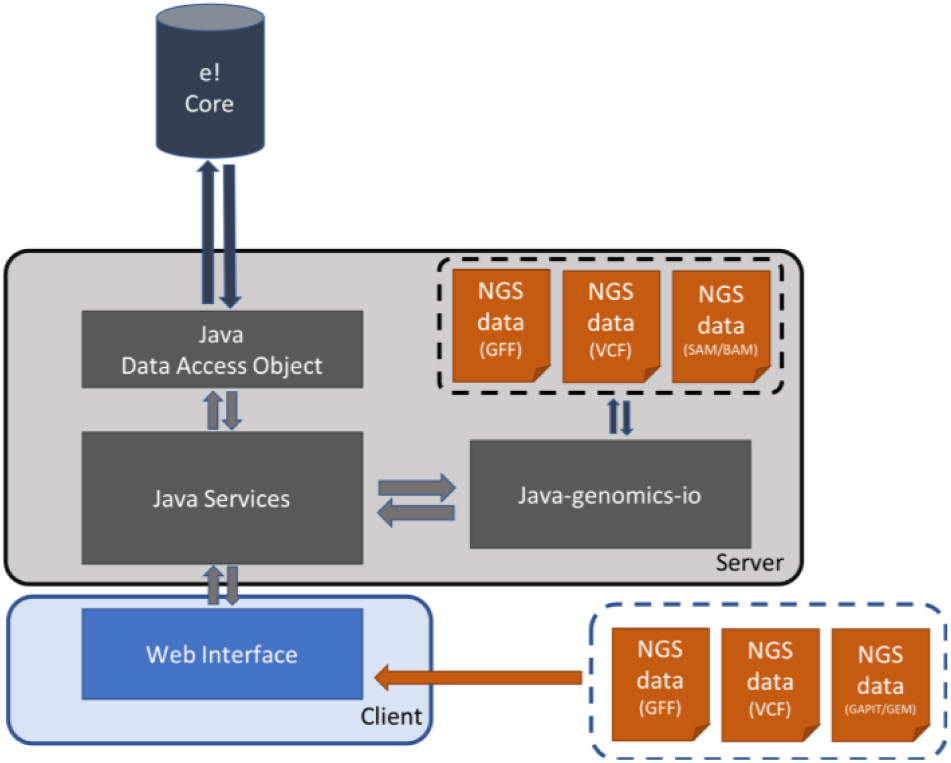
The TGAC Browser infrastructure, showing the interactions between the server-side implementation, connected to Ensembl core database using Java Data Access Objects and NGS files via Java-genomics-io, and the client-side implemented using popular techniques such as JavaScript, jQuery, d3.js and jQuery DataTables.

The server side of TGAC Browser is implemented in Java programming language, which retrieves data from a local Ensembl Core database using Java DAO (Data Access Objects) and NGS formatted files using Java-Genomics-IO library (10), a Java library developed by Timothy Palpant. TGAC Browser retrieves references from Ensembl database and visualises genomic annotation from the Ensembl database as well as NGS formatted files such as SAM (Sequence Alignment/Map format) (11), BAM (Binary equivalent of SAM), GFF (Generic Feature Format) (12), and VCF (Variant Call Format) (13).

TGAC Browser client-side is implemented in JavaScript, jQuery library, SVG (Scalable Vector Graphics) and D3.js (Data-Driven Documents) (14). By using all these well-known web technologies, we are able to create seamless browsing experience for users, where user can drag, pan and rearrange genomics tracks on a web browser similar to Google Maps. We have implemented lazy loading method, in which TGAC Browser retrieves and visualise data only for the visible and surrounding regions and for any action by the user it retrieves only required additional data. This strategy allows for very dynamic zooming and scrolling to provide smoother and faster users experience.

TGAC Browser allows users to upload (Figure 3E) genomic annotations such as GFF, GAPIT (Genome Association and Prediction Integrated Tool) (15) and GEM file format containing information about Genes, SNPs and expression data. This provides a collaborative platform for users to visualise their data safely without needing to share or making it available from the server.

**Fig. 2.**
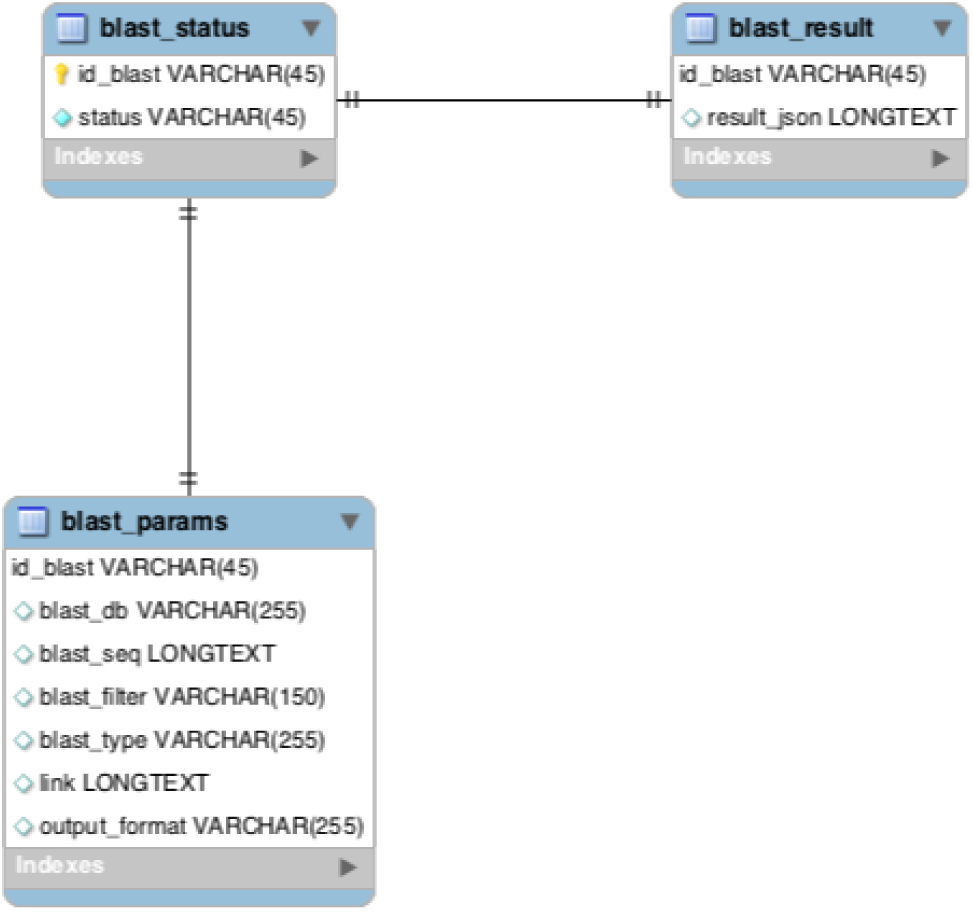
blast_manager database schema.

**Fig. 3.**
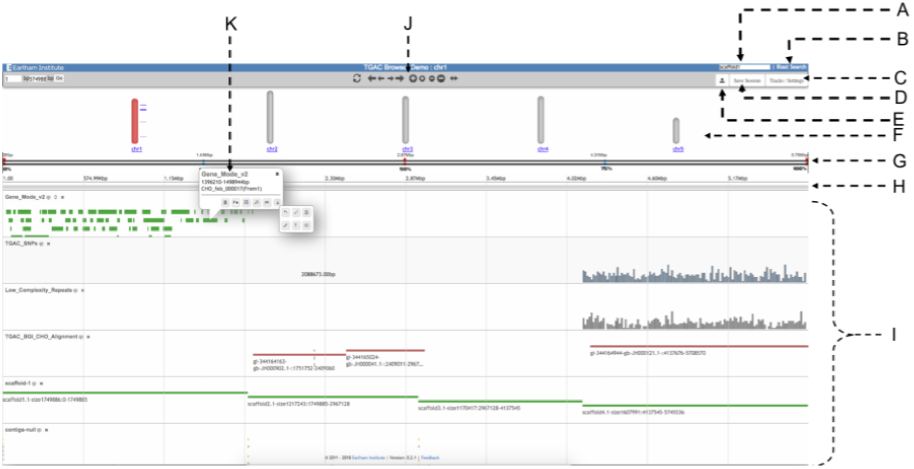
The main view of TGAC Browser. The header on top provides a search box (A) and a link to BLAST Search (B). It is followed by the second panel containing Control bar (J), an option to toggle tracks (C), save session (D) and upload tracks (D). Chromosomal view (F) represents available chromosomes for the species, where the selected chromosome is coloured in red. Below there is a horizontal view of reference (G), followed by a zoomed area of the reference (H). All genomic tracks (I) are laid out in order after that. The figure is also showing an example of a typical popup (K)

TGAC Browser has an integrated BLAST search functionality to add analysis capability. BLAST can be set up to run on local installation or High-performance computing (HPC) cluster using BLAST+ (16), as well as NCBI BLAST server. TGAC Browser keeps track of BLAST analysis using *blast_manager*, a database system (see Figure 2), and stores result for future reference.

## Results

The layout of the TGAC Browser (see Figure 3) is similar to the many of the genome browser available, making it userfriendly. In this layout, the genomic region spans from left to right and genomic annotations are laid out from top to bottom.

TGAC Browser visualises reference level genomic information on top using chromosome information (Figure 3 F), if available. as well as horizontal selectable region (Figure 3 G). User can move selector on the selectable region, and the respective region will be shown below (Figure 3 H), which can be visualised as nucleotide sequences and three forward frame translation if zoomed enough. The chromosomal view gives user overview of the reference species as well as provide a visual guide of the current viewing region on the chromosome and let user change region of interest. Chromosomal view also visualises available genomic markers on the side.

### Interface features

TGAC Browser has implemented various browsing functionalities to provide a seamless browsing experience. Navigation controls are in the top control bar (Figure 3 J) for panning and zooming. It also contains an expand button for an overview of the whole reference and reset button to focus on the centre point of reference. In addition, TGAC Browser also equipped with google maps style panning by dragging the mouse and zooming with a scroll or double click as well as panning with arrow keys on the keyboard.

Genomic annotations can be ordered by dragging them with a label of the track and toggled from Tracks/Settings (Figure 3 C). Primary information for each annotation is visualised next to the genomic track, and additional information can be seen with mouseover, as well as in a popup (detailed below).

### Search

TGAC Browser is equipped with flexible keyword-based search functionality (Figure 3 A), which searches against chromosome names, assembly information as well as all the relevant genomic features information such as gene symbols, Ensembl stable IDs (unique identifiers in the Ensembl project for each genomic annotation), common names in the database. It visualises results along with Chromosomal view if available or in tabular form with a link to respective browser view.

### Visualisations

TGAC Browser presents genomic annotations using various types of visualisations automatically chosen by the type and volume of genomic data to be visualised (see Figure 4). For small dataset each annotation visualises independently while large dataset visualises either as a histogram (Figure 4 B) or a heat map (Figure 4 A) representing quantitative information. This method is memory efficient as well as helps the user to look at a glance for a larger region and then focus on a particular segment for a detailed view. TGAC Browser also uses wiggle plots (Figure 4 C) for expression data using wigExplorer (17) from BioJS (18) and Manhattan style (Figure 4 D) visuals for SNPs.

**Fig. 4.**
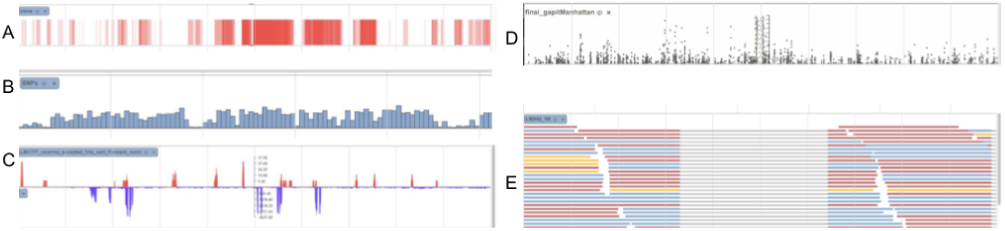
TGAC Browser visualises genomic annotation based on type and amount of data: A: Heat Map presentation of large data (more than 5000 annotation), B: Graphical presentation of large data (from 1000 to 5000 annotation), C: Wiggle plot for expression data using wigExplorer, D: Manhattan plot for GAPIT data, E: Visualising reads directly from SAM/BAM file

### Pop-up

TGAC Browser provides a contextual menu system via interactive pop-ups (Figure 3 K), which contains additional information for genomic annotation such as analysis type, position on the reference and textual description. Pop-up also contains options to fetch sequence, perform BLAST analysis for the sequence of selected annotation, focus on the annotation, highlighting annotation as well as provides a link to the Ensembl for more information (if the annotation is available in Ensembl). All this information and options are dynamically selected based on the type of annotation.

### BLAST

Integration of BLAST search within TGAC Browser plays a key role by providing the ability to perform analysis within. A user can utilise BLAST in two ways:

First, the user can perform BLAST search on a sequence of interest and results visualised in tabular view with links to specific result (Figure 5). User can perform multiple BLAST searches, and all the search results are shown as a selectable list, where user can toggle between results (Figure 5). In here user can also choose the type of BLAST (i.e. blastn, tblastn and blastx) as well as change parameters (e.g. scoring parameters, gap penalties and word-size). This feature gives TGAC Browser facility to search with a sequence in addition to the traditional keyword-based search.

**Fig. 5.**
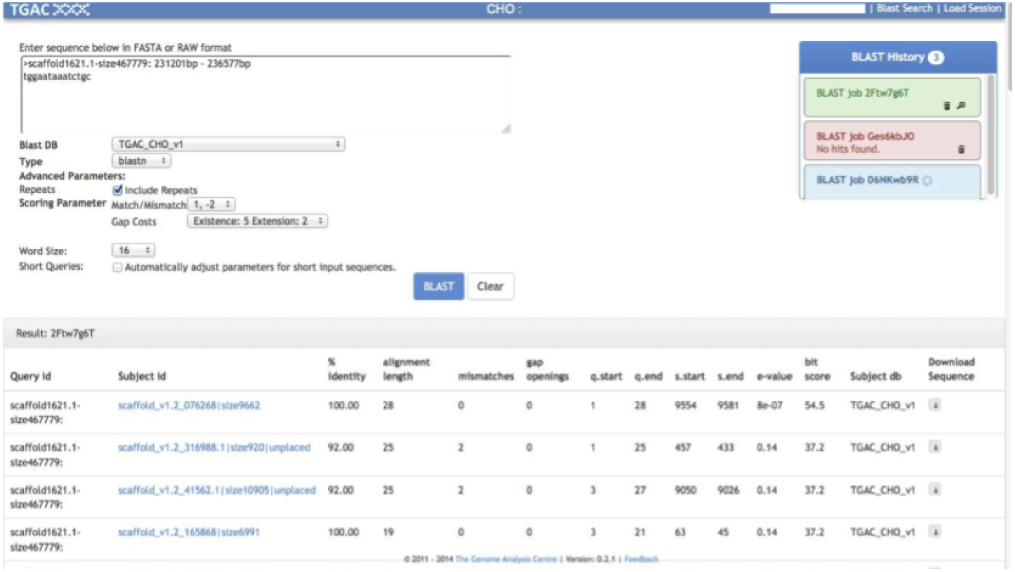
BLAST analyses are showing results with links out to associated TGAC Browser instance. on the top right showing BLAST run, allowing previous results to be shown and removed.

Second, the user can perform BLAST search from the pop-up menu of the selected genomic feature and results are presented as a genomic track alongside others (Figure 6). These BLAST Results are coloured based on standard BLAST colour schema for bit-score; it also represents insertion and deletion information. This feature helps to find out other matching regions from references for the selected annotation in the case of multiple copies of a chromosome.

**Fig. 6.**
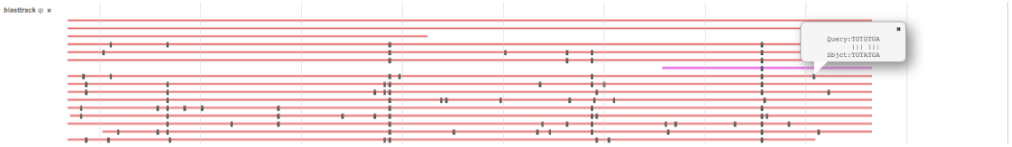
BLAST analyses are showing results as genomic track. On top right pop-up showing insertions and deletions at the position, BLAST hits are coloured based on the score.

### Sharing

TGAC Browser allows users to share information using URL as well as session id:

#### Persistent URL

TGAC Browser provides persistent unique Uniform Resource Locator (URL) to enable consistent access to the point of interest. Users can share the link for the specific reference to a given species and chromosome, or a search term. This makes it easy to share information with collaborators, or for use in publications.

#### Session

TGAC Browser also allows the user to share information using session feature (Figure 3 D), where the user can save a running session and share it with collaborator for the same view.

## Conclusions

As more and more genomes are being sequenced, genome browsers are increasing importance. Thus, we developed the TGAC Browser, a genome browser that relies on non-proprietary software but only readily available Ensembl Core database and NGS data formats. The capability of TGAC Browser to visualise data from multiple sources without any conversion makes it ideally suited to be used with newly sequenced next-generation sequencing datasets of the model and non-model organisms.

TGAC Browser follows many optimisations for data visualisations, making it versatile and robust genome browser. Functionalities to browse, upload, and share genomic information, make it all-rounder genome browser. In addition to exploration tool, TGAC Browser is also able to help scientists pursuing their research by performing analysis using built-in BLAST functionality.

TGAC Browser has been actively used by Primula Research Group at Earlham Institute and the University of Hull, SZN (Stazione Zoologica Anton Dohrn) Napoli, Brassica RIPR community, transPLANT (19), and Vietnamese Rice Community.

The ultimate goal of TGAC Browser is to provide a unique and a single solution to represent genomic data, from known NGS data format(s) for model and non-model organisms.

## Future Directions

We are looking to incorporate TGAC Browser into the virtualization system generated using CyVerse (20) and Docker with Galaxy (21) to provide a complete solution for genome analysis and exploration, where genomic annotation generated from Galaxy can directly be available to visualised using TGAC Browser instance.

We would also like to investigate into implementing, user-friendly annotation method, for users to add or modify genomic annotation. It would help to bring the community together for new genomic annotation as well as validation and curation of existing annotation.

## Availability

Information about the TGAC Browser and a demo instance are currently available at the URL below, and source code for TGAC browser is also available on GitHub. We would be pleased to help any potential users interested in the project.

Demo: http://browser.earlham.ac.uk

Source-code: https://github.com/TGAC/TGACBrowser

## ACKNOWLEDGEMENTS

This work was supported in part by the NBI Computing Infrastructure for Science Group, which provides technical support and maintenance to EI’s high-performance computing cluster and storage systems, which enabled us to develop this tool. We thank the attendees of the 2012 CHO consortium Workshop at Earlham Institute and 2012 NGS meeting at Nottingham, as well as Brassica RIPR community for helpful feedback about TGAC Browser.

## Notes

http://browser.earlham.ac.uk

## Bibliography

1. Lincoln D Stein, Christopher Mungall, Shengqiang Shu, Michael Caudy, Marco Mangone, Allen Day, Elizabeth Nickerson, Jason E Stajich, Todd W Harris, Adrian Arva, and Suzanna Lewis. The generic genome browser: a building block for a model organism system database. Genome Res., 12(10):1599–1610, October 2002.

2. W J Kent. The human genome browser at UCSC. Genome Res., 12(6):996–1006, 2002.

3. James Stalker, Brian Gibbins, Patrick Meidl, James Smith, William Spooner, Hans-Rudolf Hotz, and Antony V Cox. The ensembl web site: mechanics of a genome browser. Genome Res., 14(5):951–955, May 2004.

4. Thomas A Down, Matias Piipari, and Tim J P Hubbard. Dalliance: interactive genome viewing on the web. Bioinformatics, 27(6):889–890, March 2011.

5. Jun Wang, Lei Kong, Ge Gao, and Jingchu Luo. A brief introduction to web-based genome browsers. Brief. Bioinform., 14(2):131–143, March 2013.

6. Robert Buels, Eric Yao, Colin M Diesh, Richard D Hayes, Monica Munoz-Torres, Gregg Helt, David M Goodstein, Christine G Elsik, Suzanna E Lewis, Lincoln Stein, and Ian H Holmes. JBrowse: a dynamic web platform for genome visualization and analysis. Genome Biol., 17 (1):66, April 2016.

7. Helga Thorvaldsdóttir, James T Robinson, and Jill P Mesirov. Integrative genomics viewer (IGV): high-performance genomics data visualization and exploration. Brief. Bioinform., 14 (2):178–192, March 2013.

8. T Hubbard. The ensembl genome database project. Nucleic Acids Res., 30(1):38–41, 2002.

9. S F Altschul, W Gish, W Miller, E W Myers, and D J Lipman. Basic local alignment search tool. J. Mol. Biol., 215(3):403–410, October 1990.

10. timpalpant. timpalpant/java-genomics-io. https://github.com/timpalpant/java-genomics-io. Accessed: 2016-5-17.

11. Sequence Alignment/Map format specification. https://samtools.github.io/hts-specs/SAMv1.pdf,. Accessed: 2016-5-18.

12. GFF3 - GMOD. http://gmod.org/wiki/GFF3,. Accessed: 2016-5-18.

13. Petr Danecek, Adam Auton, Goncalo Abecasis, Cornelis A Albers, Eric Banks, Mark A DePristo, Robert E Handsaker, Gerton Lunter, Gabor T Marth, Stephen T Sherry, Gilean McVean, Richard Durbin, and 1000 Genomes Project Analysis Group. The variant call format and VCFtools. Bioinformatics, 27(15):2156–2158, August 2011.

14. Mike Bostock. D3.js - Data-Driven documents. D3. js-Data-DrivenDocuments[Internet]. [cited2015Dec21]. Availablefrom: http://d3js.org. Accessed: 2016-5-18.

15. Alexander E Lipka, Feng Tian, Qishan Wang, Jason Peiffer, Meng Li, Peter J Bradbury, Michael A Gore, Edward S Buckler, and Zhiwu Zhang. GAPIT: genome association and prediction integrated tool. Bioinformatics, 28(18):2397–2399, September 2012.

16. Christiam Camacho, George Coulouris, Vahram Avagyan, Ning Ma, Jason Papadopoulos, Kevin Bealer, and Thomas L Madden. BLAST+: architecture and applications. BMC Bioinformatics, 10:421, December 2009.

17. Anil S Thanki, Rafael C Jimenez, Gemy G Kaithakottil, Manuel Corpas, and Robert P Davey. wigexplorer, a BioJS component to visualise wig data. F1000Res., 2014.

18. John Gómez, Leyla J García, Gustavo A Salazar, Jose Villaveces, Swanand Gore, Alexander García, Maria J Martín, Guillaume Launay, Rafael Alcántara, Noemi Del-Toro, Marine Dumousseau, Sandra Orchard, Sameer Velankar, Henning Hermjakob, Chenggong Zong, Peipei Ping, Manuel Corpas, and Rafael C Jiménez. BioJS: an open source JavaScript framework for biological data visualization. Bioinformatics, 29(8):1103–1104, April 2013.

19. Manuel Spannagl, Michael Alaux, Matthias Lange, Daniel M Bolser, Kai C Bader, Thomas Letellier, Erik Kimmel, Raphael Flores, Cyril Pommier, Arnaud Kerhornou, Brandon Walts, Thomas Nussbaumer, Christoph Grabmuller, Jinbo Chen, Christian Colmsee, Sebastian Beier, Martin Mascher, Thomas Schmutzer, Daniel Arend, Anil Thanki, Ricardo Ramirez-Gonzalez, Martin Ayling, Sarah Ayling, Mario Caccamo, Klaus F X Mayer, Uwe Scholz, Delphine Steinbach, Hadi Quesneville, and Paul J Kersey. transPLANT resources for triticeae genomic data. Plant Genome, 9(1), March 2016.

20. Stephen A Goff, Matthew Vaughn, Sheldon McKay, Eric Lyons, Ann E Stapleton, Damian Gessler, Naim Matasci, Liya Wang, Matthew Hanlon, Andrew Lenards, Andy Muir, Nirav Merchant, Sonya Lowry, Stephen Mock, Matthew Helmke, Adam Kubach, Martha Narro, Nicole Hopkins, David Micklos, Uwe Hilgert, Michael Gonzales, Chris Jordan, Edwin Skidmore, Rion Dooley, John Cazes, Robert McLay, Zhenyuan Lu, Shiran Pasternak, Lars Koesterke, William H Piel, Ruth Grene, Christos Noutsos, Karla Gendler, Xin Feng, Chunlao Tang, Monica Lent, Seung-Jin Kim, Kristian Kvilekval, B S Manjunath, Val Tannen, Alexandros Stamatakis, Michael Sanderson, Stephen M Welch, Karen A Cranston, Pamela Soltis, Doug Soltis, Brian O’Meara, Cecile Ane, Tom Brutnell, Daniel J Kleibenstein, Jeffery W White, James Leebens-Mack, Michael J Donoghue, Edgar P Spalding, Todd J Vision, Christopher R Myers, David Lowenthal, Brian J Enquist, Brad Boyle, Ali Akoglu, Greg Andrews, Sudha Ram, Doreen Ware, Lincoln Stein, and Dan Stanzione. The iplant collaborative: Cyberinfrastructure for plant biology. Front. Plant Sci., 2:34, July 2011.

21. Enis Afgan, Dannon Baker, Marius van den Beek, Daniel Blankenberg, Dave Bouvier, Martin Č ech, John Chilton, Dave Clements, Nate Coraor, Carl Eberhard, Björn Grüning, Aysam Guerler, Jennifer Hillman-Jackson, Greg Von Kuster, Eric Rasche, Nicola Soranzo, Nitesh Turaga, James Taylor, Anton Nekrutenko, and Jeremy Goecks. The galaxy platform for accessible, reproducible and collaborative biomedical analyses: 2016 update. Nucleic Acids Res., 44(W1):W3–W10, July 2016.

